# Regulation of sleep quantity and intensity by long and short isoforms of SLEEPY kinase

**DOI:** 10.1101/2022.07.14.500024

**Authors:** Junjie Xu, Rui Zhou, Guodong Wang, Ying Guo, Xue Gao, Shuang Zhou, Chengyuan Ma, Lin Chen, Bihan Shi, Haiyan Wang, Fengchao Wang, Qinghua Liu

## Abstract

In *Sleepy* (*Sik3*^*Slp*^) or *Sik3*^*S551A*^ mice, deletion or mutation of inhibitory phosphorylation site serine^551^ (S551) from salt-inducible kinase 3 (SIK3) markedly increases daily non-rapid eye movement sleep (NREMS) amount, accompanied with constitutively elevated NREMS delta power density–a measure of sleep intensity or quality. Multiple SLP/SIK3 isoforms are expressed in mouse brain neurons, however, their respective roles in sleep regulation remain to be elucidated. Here, we identified a new and most abundant short isoform of SLP/SIK3 and examined sleep phenotypes resulted from isoform-specific expression of SLP-short (S) and long (L) isoforms. Adeno-associated virus-mediated adult brain chimeric (ABC)-expression of SLP-S in neurons, but not in astrocytes, significantly and constitutively elevates NREMS delta power, whereas slightly increases NREMS amount. The ability of SLP-S to regulate sleep quantity/intensity is abrogated by kinase-inactivating mutations, suggesting that the sleep-promoting activity of SLP-S is dependent on its kinase activity. In *Sik3*^*S551A-L*^ knock-in mice, isoform-specific expression of SIK3^S551A^-L (or SLP-L) significantly increases NREMS amount with a modest effect on NREMS delta power. ABC-expression of SLP-S complements the sleep phenotypes of heterozygous *Sik3*^*S551A-L*^ mice by further increasing NREMS amount and NREMS delta power to levels of *Sik3*^*Slp*^ or *Sik3*^*S551A*^ mice. Taken together, these results indicate that both SLP-L and SLP-S isoforms contribute critically to the increases of sleep quantity and intensity in *Sik3*^*Slp*^ or *Sik3*^*S551A*^ mice.

**Statement of Significance:** Previous studies have identified SIK3 as a key sleep regulatory kinase, of which gain-of-function mutations markedly increase sleep quantity and intensity in *Sleepy* mice. Multiple SLEEPY isoforms are expressed in mouse brain neurons, but their respective contributions to the hypersomnia phenotypes of *Sleepy* mice remain unclear. Here, we identified a new short isoform of SLP/SIK3 and examined sleep phenotypes resulted from specific expression of SLEEPY-short or long isoform. Our results indicate that both long and short isoforms of SLEEPY kinase contribute critically to increases of sleep quantity and intensity in *Sleepy* mice. Future studies are needed to identify downstream substrates of SLEEPY kinase that mediate regulation of sleep quantity/intensity, which should facilitate development of new treatments for human sleep disorders.

## INTRODUCTION

Sleep and wakefulness exist ubiquitously in both invertebrate and vertebrate animals. However, the molecular mechanisms that regulate sleep in mammals remain largely unclear. Previously, forward genetics screening of randomly mutagenized mice has identified a dominant *Sleepy* (*Slp*) mutation in the *Sik3* gene, which causes skipping of exon 13 in *Sik3* mRNA and in-frame deletion of 52 amino acids from SIK3 proteins [1]. The *Sleepy* (*Sik3*^*Slp*^) mice exhibit approximately 200 to 300 min increase in daily NREMS amount predominantly during the dark phase [1]. Despite of increased sleep amount, *Sik3*^*Slp*^ mice display constitutively elevated electroencephalogram (EEG) power density in 1-4 Hz (delta) frequency range during NREMS [1]. In mammals, NREMS delta power is widely used to measure sleep intensity or quality and is often regarded as one of the best indicators of sleep need [2-6].

SIK3 is an AMP-activated protein kinase (AMPK)-related protein kinase [7,8], of which the kinase activity is regulated by phosphorylation [8-10]. For example, SIK3 is activated by upstream kinases, such as LKB1, *via* phosphorylation of threonine^221^ (T221) in the activation loop of SIK3 [8,11]. On the other hand, SIK3 contains a conserved inhibitory PKA phosphorylation site, serine^551^ (S551), which is among the 52 amino acids deleted in SLEEPY (SLP) mutant proteins [1]. A non-phosphorylatable serine^551^ to alanine (S551A) substitution of SIK3 also results in marked increases of NREMS amount and NREMS delta power in *Sik3*^*S551A*^ knock-in mice [9]. Thus, mutation or deletion of S551 of SIK3 similarly creates gain-of-function mutant SLP kinase that is responsible for the hypersomnia phenotypes of *Sik3*^*S551A*^ or *Sik3*^*Slp*^ mice [9].

SIK3 proteins are predominantly and broadly expressed in the mouse brain neurons [1]. At least two SIK3/SLP isoforms, ∼145 kDa (long/L) and ∼140 kDa (medium/M), are encoded by the *Sik3/Slp* allele owning to alternative splicing [1]. Here, we identified a ∼70 KDa SIK3/SLP-short (S) isoform, which was the most abundantly expressed isoform of SIK3/SLP in the mouse brain. Because SLP-S was essentially a truncated protein lacking the carboxyl (C)-terminal half of SLP-L, we wanted to investigate the respective roles of SLP-S or SLP-L in the regulation of sleep quantity and intensity. We employed AAV-mediated adult brain chimeric (ABC)-expression system [12] for somatic genetics analysis of the sleep-promoting activity of various mutant SLP-S. Moreover, we examined the sleep phenotypes resulted from isoform-specific expression of SIK3^S551A^-L (equivalent to SLP-L) in the *Sik3*^*S551A-L*^ knock-in mice. Taken together, our results demonstrate that both SLP-L and SLP-S contribute critically to the hypersomnia phenotypes of *Sleepy* or *Sik3*^*S551A*^ mice.

## METHODS

### Animals

All animal experiments were performed according to procedures approved by the Institutional Animal Care and Use Committee of National Institute of Biological Sciences, Beijing (NIBS). All mice were provided with food and water *ad libitum*, and were housed under humidity and temperature controlled conditions (22-24□) on a 12h light: 12h dark cycle (light on at 09:00 and off at 21:00). C57BL/6J mice were purchased from the Jackson Laboratory (000664, JAX). Both *Sik3*^*Slp*^ and *Sik3*^*S551A-L*^ knock-in mice were generated at the transgenic animal facility at NIBS. The *Sik3*^*Slp*^ mice were actually *Sik3*^*E13*^∆ mice generated by deletion of exon 13 of *Sik3* gene by crossing *Sik3*^*E13-flox*^ mice with CMV-Cre mice (006054, JAX). The *Sik3*^*S551A-L*^ mice were generated by in-frame replacement of *Sik3* exon 13 genomic sequence with the *Sik3*^*S551A-L*^ (544-1369aa) open reading frame (ORF) sequence followed by the bovine growth hormone polyadenylation sequence.

### Analysis of *Sik3-S/Slp-S* ORF and plasmid construction

Mice were sacrificed by rapid cervical dislocation, and mouse brains were quickly dissected and snap-frozen in liquid nitrogen. Mouse brain tissues were ground into powder in liquid nitrogen with mortar and pestle followed by extraction of total RNA by TRIzol (Invitrogen, 15596018). Total RNA extracted from *Sik3*^*+/+*^, *Sik3*^*Slp/+*^ or *Sik3*^*Slp/Slp*^ mouse brain tissues was added to the reverse-transcriptase reaction mix for cDNA synthesis (TIAGEN, KR116-02). Forward (F: ATGGCCGC-TCGCATCGGCTA) and reverse (R: AGCACTGCCAGGTGCCAC) primers were used to amplify *Sik3-S* and/or *Slp-S* ORFs from mouse brain cDNA by PCR. The PCR products were purified by gel extraction kit (Biomed, DH101-01) and used as a template for a second PCR amplification using primers ATGGCCGCTCGCATCGGCTA and CCCGCAGATTCTGCAGTGTGC to verify the sequences encoding the N-terminal 58 amino acids of SIK3-S or SLP-S by sequencing. It should be noted, however, that the *Sik3/Slp-S* ORF that we cloned starts with the second ATG of the full-length SIK3/SLP-L isoform. We were unable to verify whether the *Sik3/Slp-S* ORF contain the first ATG by RT-PCR possibly owning to highly repetitive sequences between the first and second ATG. The pAAV-hSyn-SIK3-S or pAAV-hSyn-SLP-S was constructed by subcloning into A*ge*I-B*am*HI cleaved pAAV-hSyn-GFP (Addgene, 105539) to replace GFP with HA-tagged *Sik3-S* or *Slp-S* ORF, respectively. The pAAV-hSyn-mCherry was constructed by subcloning into A*ge*I-E*co*RI cleaved pAAV-hSyn-GFP by replacing GFP with HA-tagged mCherry ORF. The pAAV-hSyn-SLP^K37M^-S, pAAV-hSyn-SLP^T221E^-S and pAAV-hSyn-SLP^T221A^-S were derived from pAAV-hSyn-SLP-S by site-directed mutagenesis (Vazyme, P525-02). The pAAV-GFAP-SLP-S was constructed by subcloning into the S*al*I-E*co*RI cleaved pAAV-GFAP-GFP (Addgene, 50473) by replacing GFP with HA-tagged *Slp-S* ORF. The pCS2-3xFlag-SIK3-L, pCS2-3xFlag-SLP-L and pCS2-3xFlag-SLP-S were constructed by inserting *Sik3-L, Slp-L* or *Slp-S* ORF into F*se*I-A*sc*I cleaved pCS2-3xFlag-vector.

### Cell culture, transfection and immunoprecipitation

AAVpro 293T cells (Clontech, 632273) and HEK293T cells were cultured in DMEM supplemented with 10% FBS. The pCS2-3xFlag-SIK3-L, pCS2-3xFlag-SLP-L or pCS2-3xFlag-SLP-S was transfected into HEK293T cells by polyethylenimine (PEI) MAX (Polysciences, 24765). Cells were harvested 24-hour post-transfection and lysed by RIPA cell lysis buffer (Beyotime, P0013) for 20 min followed by centrifugation at 13,000 rpm for 10 min at 4ºC. The lysates (supernatants) were incubated with 30μl anti-DYKDDDDK affinity beads (Smart-Lifesciences, SA042005) for 2 hours at 4□. After washing the beads four times with 1 ml lysis buffer, the immunoprecipitants were eluted from the beads with 3xFlag peptides (MCE, HY-P0319A). The elution was mixed with SDS loading buffer and boiled at 95□ for 10 min.

### Reverse transcription and PCR (RT-PCR) analysis

Mice were sacrificed by rapid cervical dislocation. Whole mouse brain or specific sub-brain regions, such as the cortex, thalamus, hypothalamus, hippocampus, cerebellum or brain stem, were quickly dissected and snap-frozen in liquid nitrogen. Mouse brain tissues were ground into powder in liquid nitrogen with mortar and pestle followed by extraction of total RNA by TRIzol according to manufacturer’s protocol (Invitrogen, 15596018). Total RNA (1µg) was added to the reverse-transcriptase reaction mix for cDNA synthesis (TIAGEN, KR116-02). Primers *Sik3*-semi-F (GACGGGGCTGTGAACTTGGACAG) and *Sik3*-semi-R1 (CAGACCCGGGAACGGAAAGACT) or *Sik3*-semi-R2 (GCAGGGGAGCCTGCCCAAGGAC) were used to amplify *Sik3/Slp-L* or *Sik3/Slp-S* transcripts, respectively. After semi-quantitative PCR (23 cycles), the PCR products were resolved by agarose gel electrophoresis. Primers Q-*Sik3*-S-F/R (ACTTGTGTGGTCTCTTTGCTCA and GGGCTTCCAATGGCGCTTA), Q-*Slp*-S-F/R (GCAAGGGGATCCTACCCAT and GGAAAGGACAGTGGGAGTGG), Q-*Gapdh*-F/R (CGTCCCGTAGACAAAATGGT and TCGTTGATGGCAACAATCTC) were used for RT-qPCR analysis of endogenous *Sik3-S* mRNA and AAV-mediated expression of *Slp-S* mRNA in ABC-SLP-S mice. Primers Q-*S551A-L*-F/R (CAGAGGTCTCCTTCAGCATGG and AGATGGCTGGCAACTAGAAGG), Q-*Sik3-L*-F/R (GTGATGGCAGCCAGCATCTC and CCATGCTGAAGGAGACCTCTG) and Q-*Gapdh*-F/R were used for RT-qPCR analysis of *Sik3-L* and *Sik3*^*S551A-L*^ mRNA in heterozygous *Sik3*^*S551A-L*^ mice.

### Brain lysate preparation and immunoblotting

Mice were sacrificed by rapid cervical dislocation. Mouse brains were quickly dissected to collect whole brains or different sub-brain regions that were snap-frozen in liquid nitrogen. Mouse brain tissues were ground into powder in liquid nitrogen with mortar and pestle followed by lysis for 20 min in ice-cold RIPA buffer (Beyotime, P0013B) freshly supplemented with protease inhibitors (04693132001, Roche) and phosphatase inhibitors (4906837001, Roche). After centrifugation at ∼20,000g for 15 min at 4□, the supernatant (brain lysate) was transferred to a new microtube and stored at -80□ freezer. The protein concentrations of brain lysates were determined using the Modified BCA (bicinchoninic acid) Protein Assay Kit (C503051-0500, Sangon Biotech). Brain lysates was mixed with SDS-loading buffer and boiled at 95□ for 10 min. Immunoblotting was performed with the following antibodies: Anti-HA (AF0039, Beyotime), anti-β-ACTIN (AF003, Beyotime), anti-GAPDH (60004-1-Ig, Proteintech), anti-FLAG M2 HRP (Sigma, A8592), anti-14-3-3 (Santa Cruz, sc-23957) antibodies were purchased from commercial sources. Rabbit polyclonal anti-SIK3 antibodies were generated by Abcam custom antibody production service using a peptide antigen located within N-terminal 200-400 amino acids of SIK3 proteins. The specificity of the anti-SIK3 antibodies was validated by immunoblotting of whole brain lysates of wild type and ABC-*Sik3*^*KO*^ mice generated by triple-target CRISPR/Cas9 [12].

### AAV packaging and purification

Recombinant AAV-PHP.eB [13] was packaged in AAVpro 293T cells by co-transfection of PHP.eB (Addgene, 103005), pHelper (Agilent, 240071-54) and transfer plasmids using polyethylenimine (PEI) MAX (Polysciences, 24765). Cells were harvested by cell lifter (Biologix, 70-2180) after 72-hour post-transfection. Cell pellets were resuspended in 1x Gradient Buffer (10 mM Tris-HCl pH=7.6, 150 mM NaCl, 10 mM MgCl_2_). Cells were lysed by five cycles of freezing in liquid nitrogen for 5 min, thawing in 37□ water bath for 2 min and vortexing for 2 min. Then ≥50 U/mL of Benzonase nuclease (Millipore, E1014) were added to cell lysates and incubated at 37□ for 30 min. After centrifugation at ∼20,000g for 30 min at 4□, the supernatant was transferred to an iodixanol (Optiprep, D1556) step gradients (15%, 25%, 40% and 58%) for purification by ultracentrifugation. After centrifugation at ∼280,000g at 4□ for 4 hours, purified AAV particles were collected at the 40% iodixanol layer by a 2 ml sterile syringe (60013754, Mishawa), concentrated using Amicon filters (EMD, UFC801096), and formulated in sterile phosphate-buffered saline (PBS) supplemented with 0.01% Pluronic F68 (Gibco, 24040032). The titers of purified AAVs were determined by qPCR with a linearized AAV plasmid as standard.

### Immunofluorescence

Mice were completely anesthetized and subjected to cardiac perfusion with normal saline followed by 4% paraformaldehyde in PBS. Mouse brains were carefully dissected out and post-fixed in 4% paraformaldehyde in PBS at room temperature (RT) for 4 hours. After incubating in 30% sucrose for 24 hours at room temperature, mouse brains were sectioned at 35-40 micron on a cryostat microtome (Leica) under the protection of tissue OCT-Freeze medium. Brain sections were washed in PBS to remove tissue embedding agent, and followed by washing three times in PBST (0.3% Triton X-100 in PBS) for 5 min at RT. After incubating in the blocking solution (3% BSA in PBST) for 1 hour at RT, brain sections were incubated with the primary antibodies in blocking solution overnight at 4□. The following primary antibodies were used: anti-HA (1:500, 11867423001, Roche), anti-NeuN (1:500, ABN78, Milipore), anti-S100β (1:500, ab52642, Abcam). After washing three times in PBS for 5 min at RT, brain sections were incubated with fluorescence-tagged secondary antibodies in PBS for 2 hours at RT. The following secondary antibodies were used: Alexa Fluor 555 conjugated goat-anti-rat (1:500, 4417, CST), Alexa Fluor 488 conjugated goat-anti-rabbit (1:500, 111-545-003, Jackson Immunoresearch). After washing three times in PBS, the brain sections were mounted on adhesion microscope slides (Genview) and encapsulated in sealed tablets containing 3 μg/ml DAPI solution (C0060, Solarbio). Fluorescent images were obtained using Digital Slicing Workstation (VS120, Olympus) and Ultra-high resolution laser confocal microscope (LSM800, Zeiss). For quantification of AAV transduction rates, 8 to 10 different windows were randomly selected for counting in the cortex, thalamus or hypothalamus from immunostaining images of the same mouse. Imaris 7.6.0 and ImageJ were used to process and count the number of AAV-transduced neurons or astrocytes in these images.

### EEG/ encephalogram (EMG)-based sleep analysis

For EEG/EMG recording of sleep/wake behaviors, 12 to 13 weeks old adult male mice were anesthetized by isoflurane (4% for induction, 1% for maintenance). After confirming that the mice lost consciousness, the head region was shaved, and the skull was exposed with sterilized surgical tools. Cotton swabs with hydrogen peroxide was used to clean the exposed skull to improve adhesion of skull and dental cement. After the coordinate of lambda point was set as (0, 0, 0), four coordinates (−1.27, 0, 0), (−1.27, 5.03, 0), (1.27, 5.03, 0) and (1.27, 0, 0) were sequentially set as the insertion sites for four electrode pins. Hand-held electric drill was used to punch holes in the skull and EEG electrode pins were implanted to the dura at a depth of 1.3 mm under the skull surface by stereotaxic control. EEG pins at insertion sites (±1.27, 5.03,1.3) targeted to motor cortex M1, and at (±1.27, 0,1.3) targeted to secondary visual cortex V2ML. Two EMG wires were inserted into the neck muscle and the EEG electrode in the skull was then fixed with dental cement. After surgery, the mice were singly housed to recover for at least 24 hours. Then, 200μl recombinant AAVs were retro-orbitally injected into mice (10^12^ vg/mouse) in one week after surgery. EEG/EMG recording normally started after two weeks post AAV injection.

After acclimation in the recording room for one week (included in the two-week period between AAV injection and EEG/EMG recording), mice are subjected to EEG/EMG recording for three consecutive days. The sleep/wake behaviors were analyzed as previously described with slight modifications [1,14]. In brief, EEG/EMG recording data are visualized and analyzed using a custom-designed, C++ language-based semi-automated sleep/wake staging software: 1) EEG signals are subjected to fast Fourier transform analysis for 1 to 30 Hz with 1 Hz bins; 2) Our software divides each 20-s epoch into five 4-s epoch and stages sleep/wake states in a 4-s epoch: NREMS [high amplitude of delta frequency (1–4 Hz) EEG and low EMG tonus], rapid eye movement sleep (REMS) [high amplitude of theta frequency (6–9Hz) EEG and EMG muscle atonia] and Wake (low amplitude, fast EEG and high amplitude, variable EMG signal); 3) Our software applies a “smooth” process to every 20-s epoch to classify it as NREMS, REMS or Wake state based on the context; 4) Finally, the results of automated sleep/wake staging are manually inspected and corrected. The relative NREMS delta power density was calculated by the ratio of delta power (1–4 Hz) to total power (1–30 Hz) of EEG signals during NREMS. The hourly plot of NREMS delta power was calculated by averaging the relative NREMS delta power density in each hour during the three-day recording period. The EEG power spectra analysis was conducted by calculating the ratio of each 1 Hz bin to total 1–30 Hz of EEG signals in NREMS during the 24-h cycle. While “episode number” was the average number of NREMS, REMS or Wake state during the 24-h cycle, “episode duration” was the mean episode duration of NREMS, REMS or Wake state during the 24-h cycle [zeitgeber time (ZT)0-ZT24], light phase (ZT0-ZT12) or dark phase (ZT12-ZT24).

### Statistical analysis

No statistical methods were used to predetermine sample size. Randomization and blinding were not used except that the technicians who performed EEG/EMG-based sleep analysis were blind to the genotypes of mice or the types of AAVs injected. All relevant data were shown as mean ± s.e.m. GraphPad Prism 9.0.0. was used for the quantitative analysis or the statistical analysis of the NREMS, REMS and wake data. Typically, unpaired t test was used for pairwise comparisons; one-way ANOVA was used for multiple comparisons; two-way ANOVA was applied for multiple comparisons involving two independent variables. Pearson’s chi-squared test was applied to determine whether there is a statistically significant difference between the expected and observed frequencies for homozygous *Sik3*^*S551A-L*^ mice. The * *p* < 0.05, ** *p* < 0.01, *** *p* < 0.001 and **** *p* < 0.0001 levels were considered statistically significant.

## RESULTS

### Identification and characterization of a new SIK3/SLP-S isoform

In addition to two previously reported ∼145 kDa and ∼140 kDa SIK3/SLP isoforms [1], we detected a new ∼70 kDa short isoform of SIK3/SLP by immunoblotting of whole brain lysates from *Sik3*^*+/+*^, *Sik3*^*Slp/+*^ and *Sik3*^*Slp/Slp*^ mice using the amino (N)-terminal specific anti-SIK3 antibodies **(Figure 1, A-C and Supplementary Figure S1A)**. A corresponding SIK3-S isoform is annotated in the genome database (UniProt F6U8X4), which lacks the N-terminal 58 amino acids, including lysine^37^ (K37) that is required for the kinase activity of SIK3 [11]. Thus, we cloned the endogenous *Sik3-S* or *Slp-S* cDNA from *Sik3*^*Slp/+*^ mouse brain by reverse transcription (RT)-PCR and verified by sequencing that SIK3-S or SLP-S included the N-terminal 58 amino acids **(Figure 1D and Supplementary Figure S1B)**. As annotated in the database, both *Sik3-S* and *Slp-S* transcripts terminate after exon 14 with the use of alternative polyadenylation sequence **(Figure 1, A and B)**. Because of a premature stop codon right after exon 14, SIK3/SLP-S is essentially a truncated protein lacking the C-terminal half portion of SIK3/SLP-L/M. Consistent with a previous report [9], we showed that both SLP-L and SLP-S showed reduced binding of 14-3-3 proteins relative to SIK3-L by co-immunoprecipitation in transfected 293T cells **(Supplementary Figure S1C)**.

**Figure 1.**
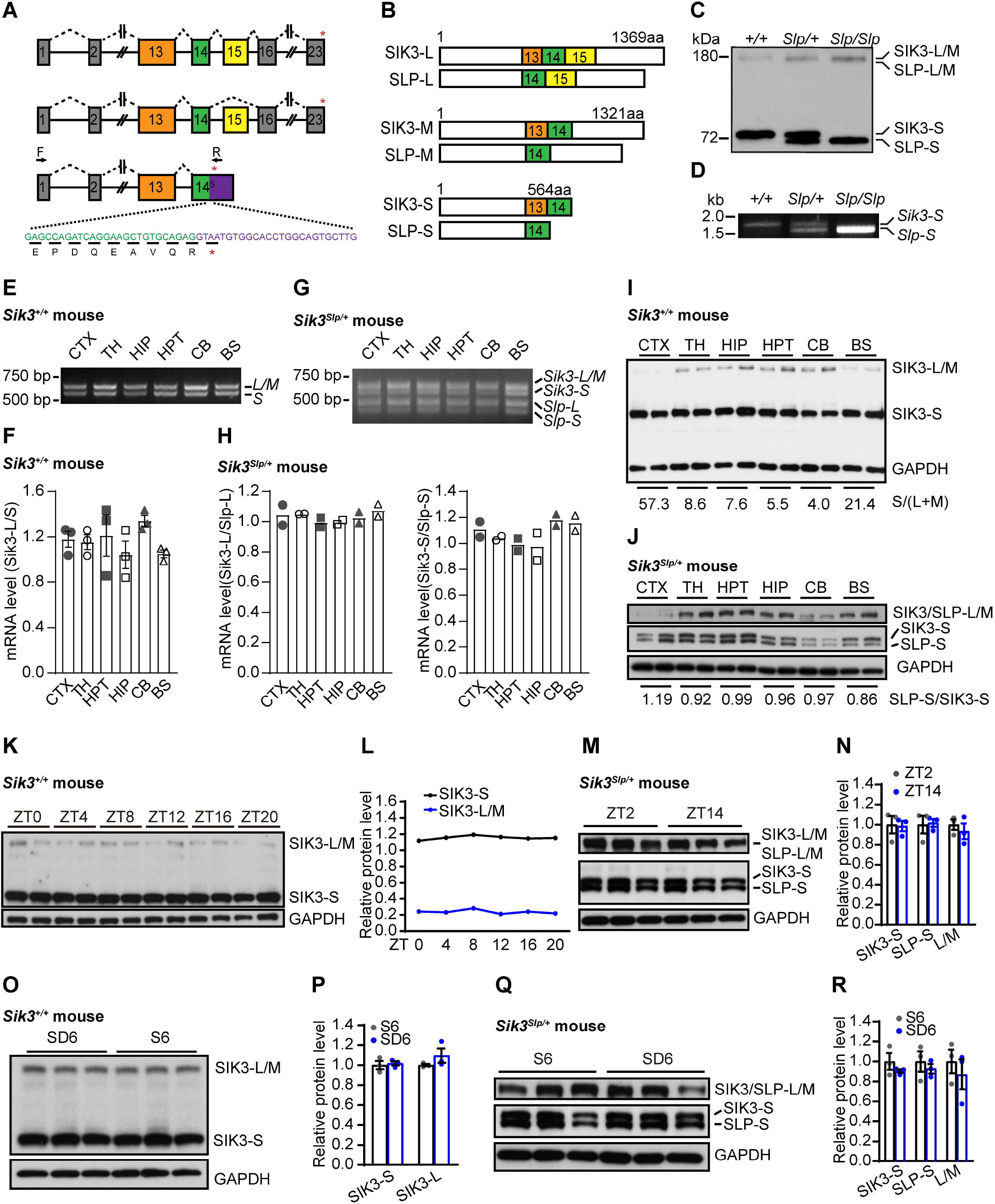
Identification and characterization of a new SIK3/SLP-S isoform. **(A and B)** Schematic of different isoforms of *Sik3* mRNA (A) and SIK3 proteins (B) owning to alternative splicing or polyadenylation usage. Boxes represent exons and lines as introns. F/R refer to primers for RT-PCR analysis of *Sik3-S* mRNA. Red star indicates stop codon. **(C)** Immunoblotting of *Sik3*^*+/+*^, *Sik3*^*Slp/+*^ or *Sik3*^*Slp/Slp*^ mouse brain lysates using anti-SIK3 antibody. **(D)** Analysis of *Sik3-S* or *Slp-S* mRNA by RT-PCR in *Sik3*^*+/+*^, *Sik3*^*Slp/+*^ and *Sik3*^*Slp/Slp*^ mouse brains. **(E)** Semi-quantitative RT-PCR analysis of *Sik3-L* and *Sik3-S* transcripts in six brain regions of C57BL/6J mouse. CTX, cortex; TH, thalamus; HIP, hippocampus; HPT, hypothalamus; CB, cerebellum; BS, brain stem. **(F)** Quantitation of the ratio of *Sik3-L/Sik3-S* transcripts in (D) (n=3). **G)** Semi-quantitative RT-PCR analysis of *Sik3/Slp-L and Sik3/Slp-S* transcripts in different brain regions of *Sik3*^*Slp/+*^ mouse. **(H)** Mean ratio of *Sik3-L/Slp-L* or *Sik3-S/Slp-S* transcripts in (F) (n=2). **(I)** Immunoblotting of lysates of different brain regions of C57BL/6J mouse. The ratio of SIK3-S/SIK3-(L/M) proteins (normalized to GAPDH) is listed below. **(J)** Immunoblotting of lysates of different brain regions of *Sik3*^*Slp/+*^ mouse. The ratio of SLP-S/SIK-S (normalized by GAPDH) is listed below. **(K)** Immunoblotting of whole brain lysates of C57BL/6J mice at different ZT times. **(L)** 24-h plot of mean level (n=2) of SIK3-L/M and SIK3-S proteins (normalized to GAPDH) in (J). **(M)** Immunoblotting of whole brain lysates of *Sik3*^*Slp/+*^ mice at ZT2 and ZT14. **(N)** Quantitation of the levels of SIK3-S or SLP-S proteins between ZT2 and ZT14. Two-way ANOVA with Sidak’s test: F (2, 12) =0.196. Data are mean ± s.e.m. Not significant (*p*>0.05) is not shown. **(O)** Immunoblotting of whole brain lysates of C57BL/6J mice after 6-h sleep deprivation (SD6) or *ad libitum* sleep (S6). **(P)** Quantitation of the levels of SIK3-L/M or SIK3-S proteins in (N). Two-way ANOVA with Sidak’s test: F (1, 8) =0.799. Data are mean ± s.e.m. Not significant (*p*>0.05) is not shown. **(Q)** Immunoblotting of whole brain lysates of *Sik3*^*Slp/+*^ mice after SD6 and S6 treatments. **(R)** Quantitation of the levels of SIK3-L/M, SIK3-S or SLP-S proteins in (P). Two-way ANOVA with Sidak’s test: F (2, 12) =0.044. Data are mean ± s.e.m. Not significant (*p*>0.05) is not shown.

To study whether *Sik3/Slp* isoforms are differentially expressed in different brain regions, we compared the mRNA/protein levels of different *Sik3/Slp* isoforms in the cortex, thalamus, hypothalamus, hippocampus, brain stem and cerebellum of wild-type and *Sik3*^*Slp/+*^ mice. As shown by semi-quantitative RT-PCR, there were similar levels of *Sik3-L* and *Sik3-S* mRNA in all six brain regions of wild-type mice, whereas the level of *Slp-L*/S mRNA was comparable to that of *Sik3-L/S* mRNA **(Figure 1, E-H)**. In contrast, the relative ratio of SIK3-S vs. SIK3-L/M proteins varied among different brain regions, ranging from the lowest (∼4:1) in the cerebellum to the highest (∼57.3:1) in the cortex of wild-type mice **(Figure 1I)**. Moreover, the level of SLP-S was largely comparable to that of SIK3-S among all six brain regions in *Sik3*^*Slp/+*^ mice **(Figure 1J)**.

To examine whether the expression of SI K3 isoforms was regulated by circadian rhythm, we collected brain samples from wild-type mice at every 4-h during the 24-h cycle. Immunoblotting showed that the levels of all SIK3 isoforms did not change significantly during the circadian cycle **(Figure 1, K and L)**, which was different from a recent report of circadian fluctuation of SIK3 proteins [15]. Likewise, there was no significant difference in the expression of SIK3/SLP isoforms in *Sik3*^*Slp/+*^ mouse brains between ZT2 (light phase) and ZT14 (dark phase) **(Figure 1, M and N)**. Neither did we observe any significant change of SIK3/SIK3 isoforms in wild-type or *Sik3*^*Slp/+*^ mice after 6-h sleep deprivation **(Figure 1, O-R)**. It should be noted, however, that we could not exclude the possibility of significant change in the expression of SIK3/SLP isoforms in specific sub-brain regions during the diurnal cycle or after sleep deprivation.

### ABC-expression of SLP-S in neurons significantly elevates NREMS delta power, whereas slightly increases NREMS amount

Retro-orbital injection of AAV-PHP.eB, a mutant serotype derived from AAV9, could bypass the blood brain barrier and efficiently transduce the majority of neurons and astrocytes across the adult mouse brain [12,13]. Importantly, AAV-PHP.eB-mediated systemic expression of GFP reporter across the mouse brains results in no alteration of the sleep-wake architecture [12]. To study sleep regulatory functions of SIK3/SLP-S, we performed ABC-expression of GFP, SIK3-S or SLP-S from human synapsin (hSyn) promoter in the adult brain neurons of C57BL/6J mice by retro-orbital injection of AAV-hSyn-GFP, AAV-hSyn-SIK3-S or AAV-hSyn-SLP-S [12,16,17]. Co-immunostaining of HA and NeuN, a pan-neuronal marker protein, indicated systemic expression of HA-tagged GFP, SIK3-S or SLP-S in ∼50 to 80% of neurons in the cortex, thalamus and hypothalamus of AAV-injected mice **(Supplementary Figure S2, A-D)**. While there were similar levels of AAV-derived *Sik3-S* and *Slp-S* transcripts, the level of SIK3-S proteins was higher than that of SLP-S proteins as shown by immunoblotting **(Supplementary Figure S2, E and F)**. Moreover, the level of AAV-mediated expression of SLP-S was ∼30% of that of endogenous SIK3-S proteins in the ABC-SLP-S mouse brain lysates **(Supplementary Figure S2, G and H)**.

Whereas ABC-expression of SIK3-S did not alter the sleep-wake architecture, ABC-expression of SLP-S resulted in a moderate sleepy phenotype **(Figure 2)**, consistent with that SLP-S was a gain-of-function mutant of SIK3-S. Specifically, ABC-SLP-S mice, relative to ABC-GFP control mice, showed modest (average ∼55 min/day) increase of NREMS time during the dark phase, reduction of REMS time during the light phase, and decrease of wake time during the dark phase **(Figure 2, A and B)**. Moreover, ABC-SLP-S mice showed less NREMS episode number and longer NREMS episode duration, indicative of more consolidated NREMS **(Figure 2, C and D)**. Importantly, ABC-SLP-S mice displayed significantly and constitutively elevated (by ∼27.9%) NREMS delta power relative to ABC-SIK3-S and ABC-GFP mice **(Figure 2, E and F)**. There was also significant reduction of theta power density during NREMS as well as modest increase of delta power density during REMS and wakefulness **(Figure 2F)**. These results suggest that ABC-expression of SLP-S in neurons significantly elevates NREMS delta power, whereas slightly increases NREMS amount.

**Figure 2.**
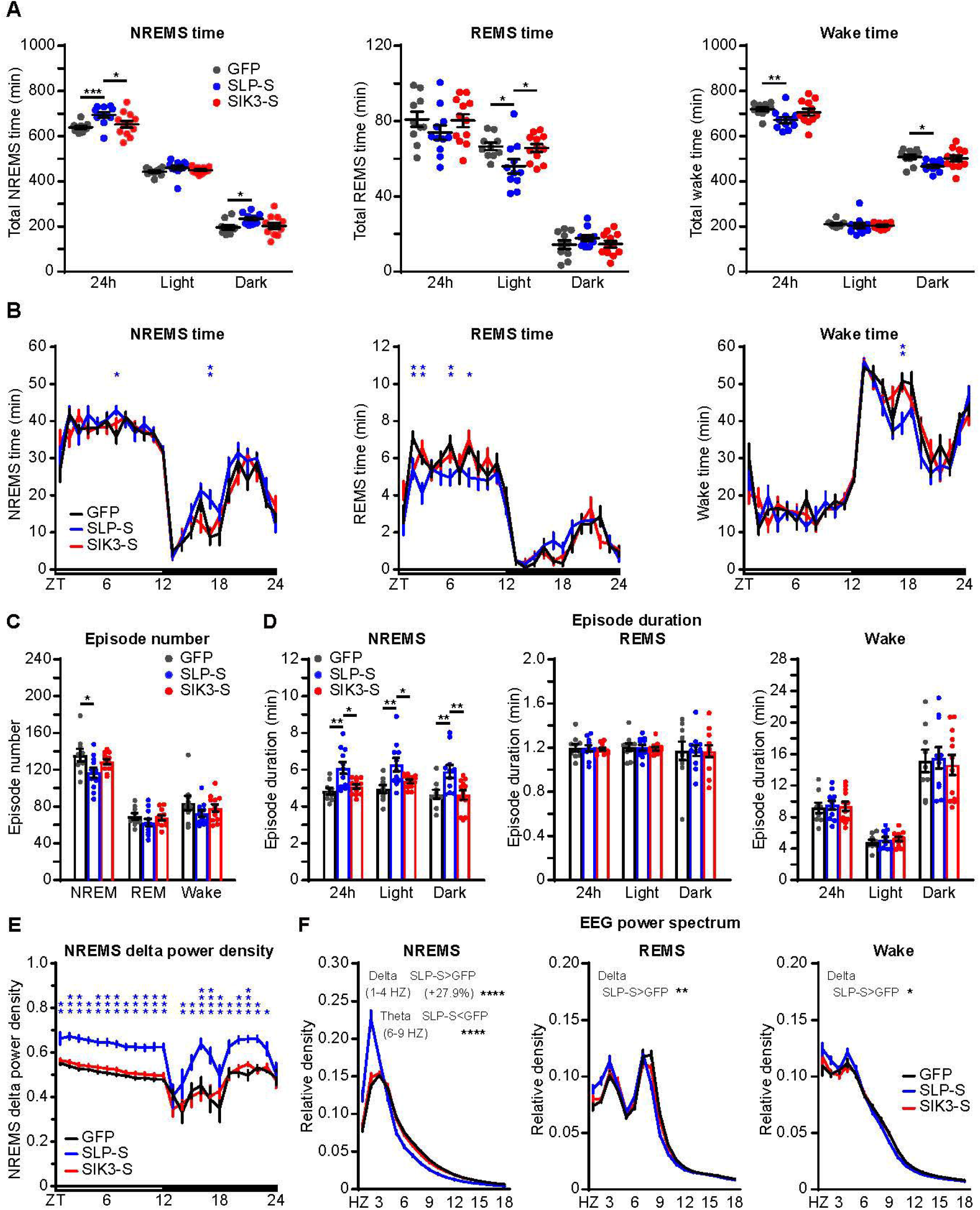
ABC-expression of SLP-S in neurons significantly elevates NREMS delta power and slightly increases NREMS amount. **(A)** Quantification of daily NREMS, REMS or wake time in C57BL/6J adult mice retro-orbitally injected with AAV-hSyn-GFP (ABC-GFP, n=10), AAV-hSyn-SLP-S (ABC-SLP-S, n=11), or AAV-hSyn-SIK3-S (ABC-SIK3-S, n=12). Two-way ANOVA with Tukey’s test: NREMS, F (4, 90) =1.026; REMS, F (4, 90) =1.979; Wake, F (4, 90) =1.330. **(B)** Hourly plots of NREMS, REMS or Wake time in the ABC-GFP, ABC-SLP-S and ABC-SIK3-S mice. Two-way ANOVA with Dunnett’s test: NREMS, F (46, 720) =1.612; REMS, F (46, 720) =2.303; Wake, F (46, 720) =1.781. **(C and D)** Quantification of NREMS, REMS and Wake episode numbers (C) or episode durations (D) in the ABC-GFP, ABC-SLP-S and ABC-SIK3-S mice. Two-way ANOVA with Tukey’s test: Episode number, F (4, 90) =0.464; NREMS episode duration, F (4, 90) =0.250; REMS episode duration, F (4, 90) =0.007; Wake episode duration, F (4, 90) =0.114. **(E)** Hourly plot of relative NREMS delta power density in the ABC-GFP, ABC-SLP-S and ABC-SIK3-S mice. Two-way ANOVA with Dunnett’s test: F (46, 720) =0.981. **(F)** EEG power spectra analysis of NREMS, REMS and Wake states (ZT0-ZT24) in the ABC-GFP, ABC-SLP-S and ABC-SIK3-S mice. One-way ANOVA with Tukey’s test. Delta (1-4 Hz): NREMS, F (2, 30) =51; REMS, F (2, 30) =8.205; Wake, F (2, 30) =5.454. Theta (6-9 Hz): NREMS, F (2, 30) =50.370; REMS, F (2, 30) =2.172; Wake, F (2, 30) =1.114. Data are mean ± s.e.m. * *p*<0.05; ** *p*<0.01; *** *p*<0.001; **** *p*<0.0001. Not significant (*p*>0.05) is not shown.

### ABC-expression of SLP-S in astrocytes does not cause sleepy phenotypes

Next, we performed ABC-expression of SLP-S in astrocytes by retro-orbital injection of C57BL/6J adult mice with AAV-PHP.eB expressing GFP or SLP-S from the glial fibrillary acidic protein (GFAP) promoter, which could deliver expression of target gene in most if not all types of astrocytes in the mouse brain [18]. Co-immunostaining of HA and S100□, an astrocyte-specific maker protein, indicated systemic expression of HA-tagged GFP or SLP-S in ∼60% to 80% of astrocytes of these AAV-injected mouse brains **(Supplementary Figure S2I)**. There was very little non-specific expression of SLP-S in neurons as shown by co-immunostaining of HA and NeuN **(Supplementary Figure S2I)**. In contrast to ABC-expression of SLP-S in neurons **(Figure 2)**, ABC-expression of SLP-S in astrocytes did not result in sleepy phenotypes, but instead cause modest reduction NREMS delta power density, increase of REMS episode duration, and EEG abnormalities **(Supplementary Figure S3)**. Because SIK3 is predominantly expressed in neurons [1,19], these minor sleep phenotypes may be attributed to inappropriate expression of SLP-S in astrocytes.

### The sleep-promoting activity of SLP-S is dependent on its kinase activity

To address whether the kinase activity of SLP-S is required for sleep regulation, we performed ABC-expression of GFP, SLP-S or SLP^K37M^-S in the adult brain neurons of C57BL/6J mice by intravenous administration of AAV-hSyn-GFP, AAV-hSyn-SLP-S or AAV-hSyn-SLP^K37M^-S **(Figure 3A)**. Notably, ABC-expression of SLP^K37M^-S, which carried a kinase-dead K37M mutation [11], caused none of the sleep phenotypes described above for ABC-expression of SLP-S, such as increases of NREMS amount and NREMS delta power **(Figure 3, B-G)**. Rather, ABC-SLP^K37M^-S mice exhibited essentially the same sleep-wake architecture as control ABC-GFP mice, except for slight reduction of wake episode number and increase of wake episode duration during the dark phase **(Figure 3, D and E)**. These results suggest that the sleep-promoting activity of SLP-S is dependent on its kinase activity.

**Figure 3.**
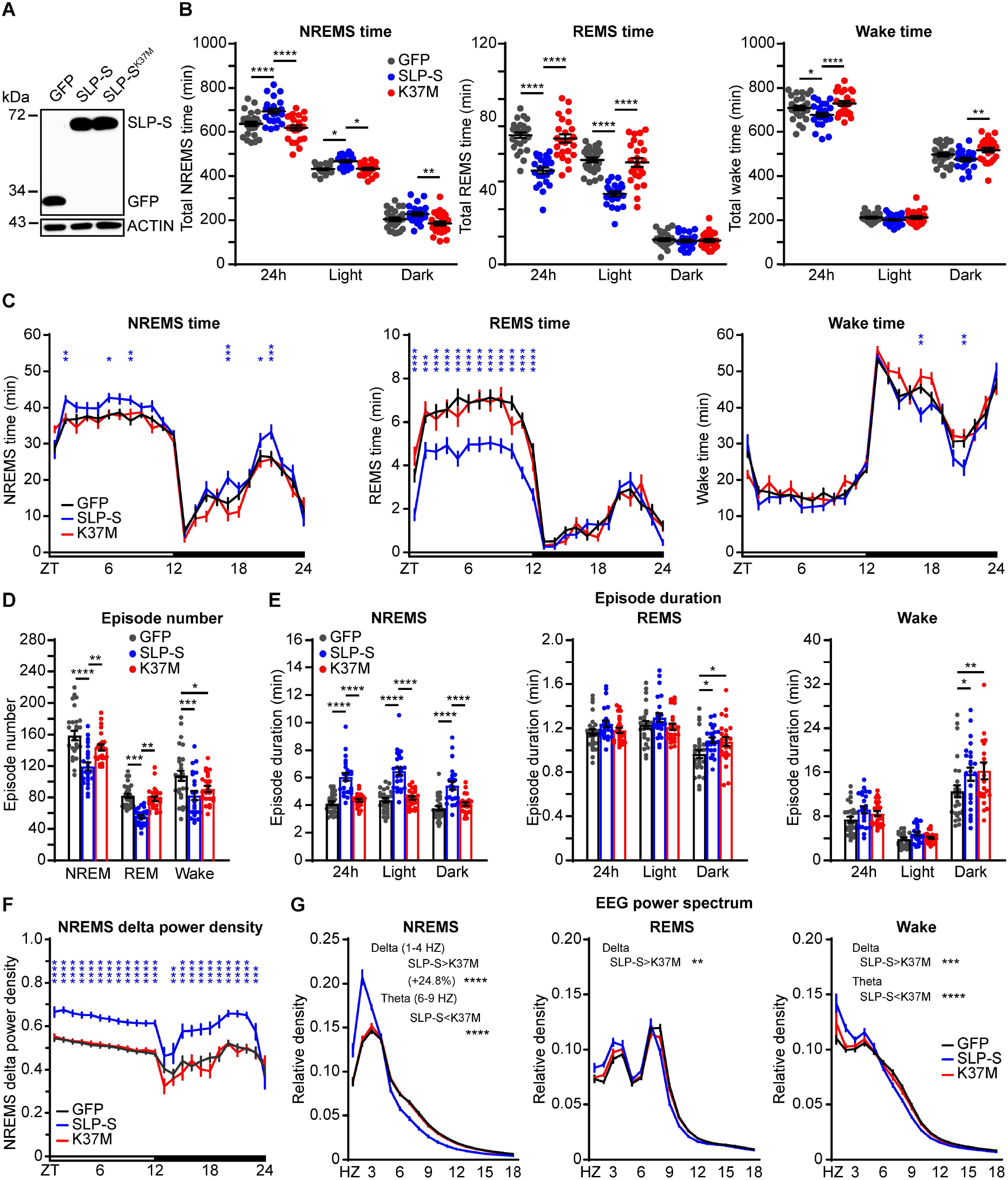
The sleep-promoting activity of SLP-S is dependent on its kinase activity. **(A)** Immunoblotting of whole brain lysates of AAV-hSyn-GFP, AAV-hSyn-SLP-S or AAV-hSyn-SLP^K37M^-S injected C57BL/6J mice. **(B)** Quantification of daily NREMS, REMS or Wake time in the ABC-GFP (n=27), ABC-SLP-S (n=23) and ABC-SLP^K37M^-S (n=24) mice. Two-way ANOVA with Tukey’s test: NREMS, F (4, 213) =2.097; REMS, F (4, 213) =14.58; Wake, F (4, 213) =1.555. **(C)** Hourly plots of NREMS, REMS or Wake time in the ABC-GFP, ABC-SLP-S and ABC-SLP^K37M^-S mice. Two-way ANOVA with Dunnett’s test: NREMS, F (46, 1704) =2.177; REMS, F (46, 1704) =4.838; Wake, F (46, 1704) =2.253. **(D and E)** Quantification of NREMS, REMS and Wake episode numbers (D) or episode durations (E) in the ABC-GFP, ABC-SLP-S and ABC-SLP^K37M^-S mice. Two-way ANOVA with Tukey’s test: Episode number, F (4, 213) =1.341; NREMS episode duration, F (4, 213) =0.646; REMS episode duration, F (4, 213) =1.115; Wake episode duration, F (4, 213) =1.338. **(F)** Hourly plot of relative NREMS delta power density in the ABC-GFP, ABC-SLP-S and ABC-SLP^K37M^-S mice. Two-way ANOVA with Dunnett’s test: F (46, 1704) =2.263. **(G)** EEG power spectra analysis of NREMS, REMS and Wake states (ZT0-ZT24) in the ABC-GFP, ABC-SLP-S and ABC-SLP^K37M^-S mice. One-way ANOVA with Tukey’s test. Delta (1-4 Hz): NREMS, F (2, 72) =74.59; REMS, F (2, 72) =13.63; Wake, F (2, 72) =18.01. Theta (6-9 Hz): NREMS, F (2, 72) =57.18; REMS, F (2, 72) =3.887; Wake, F (2, 72) =14.87. Data are mean ± s.e.m. * *p*<0.05; ** *p*<0.01; *** *p*<0.001; **** *p*<0.0001. Not significant (*p*>0.05) is not shown.

SIK3 is activated by phosphorylation of T221 [8,11,15]. Whereas phosphorylation mutation T221A diminishes, phosphorylation-mimic mutation T221E activates the kinase activity of SIK3 [11,15]. Thus, we examined the sleep-promoting activity of SLP^T221E^-S or SLP^T221A^-S in C57BL/6J adult mice by intravenous injection of AAV-hSyn-GFP, AAV-hSyn-SLP^T221E^-S or AAV-hSyn-SLP^T221A^-S **(Figure 4A)**. ABC-expression of SLP^T221E^-S caused marked (average ∼145 min) increase of daily NREMS time as well as modest reduction of REMS time during the light phase and decrease of wake time during the dark phase **(Figure 4, B and C)**. Additionally, ABC-SLP^T221E^-S mice displayed less numbers of NREMS or REMS episodes accompanied with longer NREMS or REMS episode duration, indicative of more consolidated sleep **(Figure 4, D and E)**. Moreover, ABC-SLP^T221E^-S mice exhibited significant increase of delta power and decrease of theta power during NREMS and wakefulness as well as subtle EEG alterations during REMS **(Figure 4, F and G)**. By contrast, ABC-SLP^T221A^-S mice exhibited none of these sleep phenotypes except for a slight elevation of NREMS delta power **(Figure 4, F and G)**. These observations suggest that T221 phosphorylation is critical for the sleep-promoting activity of SLP-S, which is consistent with a recent report of reduced NREMS delta power and impaired homeostatic sleep response in *Sik3*^*T221A*^ mice [15].

**Figure 4.**
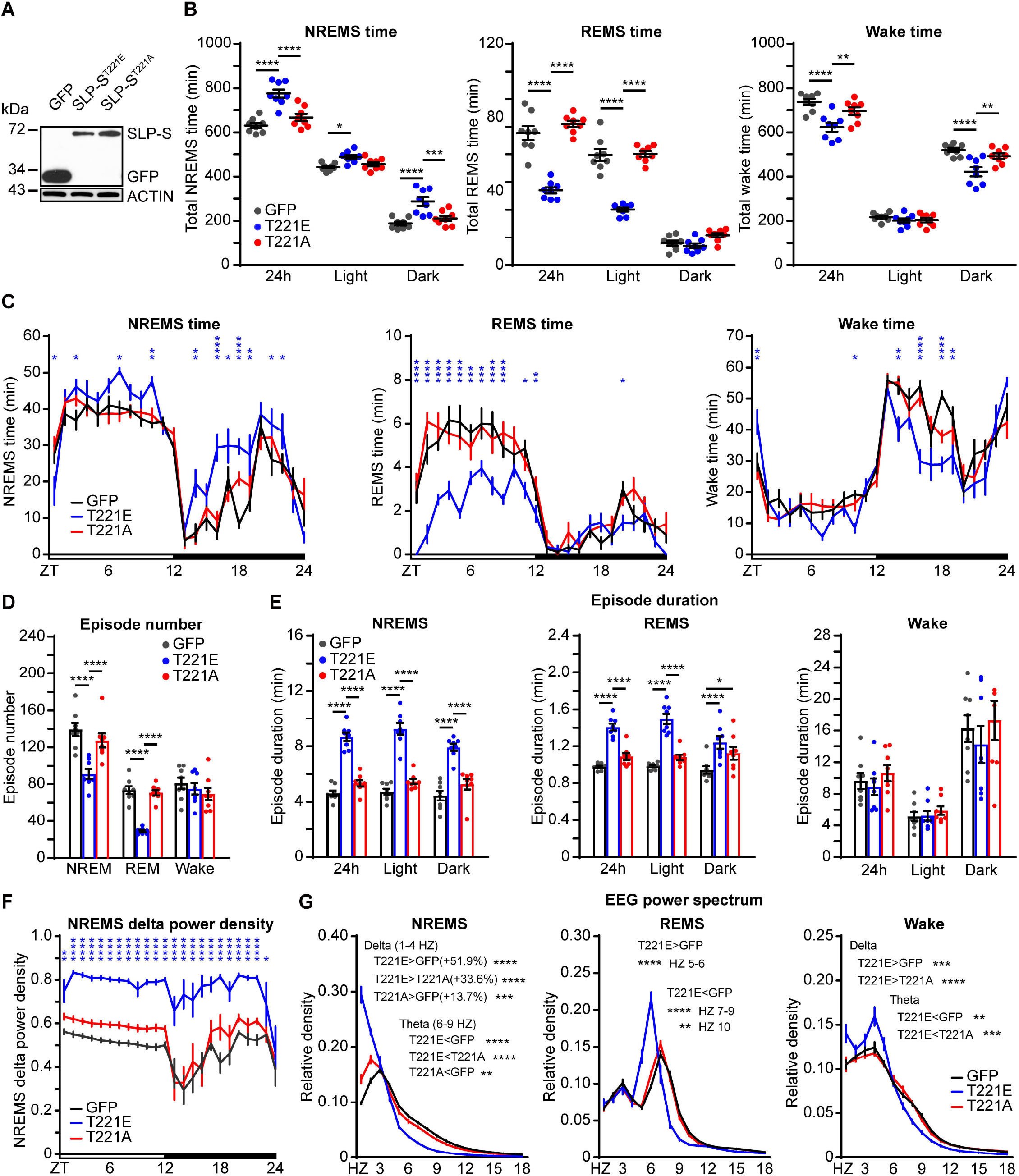
T221 phosphorylation is critical for sleep-promoting activity of SLP-S. **(A)** Immunoblotting of whole brain lysates of AAV-hSyn-GFP, AAV-hSyn-SLP^T221E^-S and AAV-hSyn-SLP^T221A^-S injected mice. **(B)** Quantification of daily NREMS, REMS or Wake time in the ABC-GFP (n=8), ABC-SLP^T221E^-S (n=8) and ABC-SLP^T221A^-S (n=8) mice. Two-way ANOVA with Tukey’s test: NREMS, F (4, 63) =3.87; REMS, F (4, 63) =21.04; Wake, F (4, 63) =3.81. **(C)** Hourly plots of NREMS, REMS or Wake time in the ABC-GFP, ABC-SLP^T221E^-S and ABC-SLP^T221A^-S mice. Two-way ANOVA with Dunnett’s test: NREMS, F (46, 504) =2.789; REMS, F (46, 504) =4.759; Wake, F (46, 504) =2.865. **(D and E)** Quantification of NREMS, REMS and Wake episode numbers (D) or episode durations (E) in the ABC-GFP, ABC-SLP^T221E^-S and ABC-SLP^T221A^-S mice. Two-way ANOVA with Tukey’s test: Episode number, F (4, 63) =6.614; NREMS episode duration, F (4, 63) =1.178; REMS episode duration, F (4, 63) =3.106; Wake episode duration, F (4, 63) =0.202. **(F)** Hourly plot of relative NREMS delta power density in the ABC-GFP, ABC-SLP^T221E^-S and ABC-SLP^T221A^-S mice. Two-way ANOVA with Dunnett’s test: F (46, 504) =1.243. **(G)** EEG power spectra analysis of NREMS, REMS and Wake states (ZT0-ZT24) in the ABC-GFP, ABC-SLP^T221E^-S and ABC-SLP^T221A^-S mice. One-way ANOVA with Tukey’s test in NREMS and Wake analysis. Delta (1-4 Hz): NREMS, F (2, 21) =172.7; Wake, F (2, 21) =18.94. Theta (6-9 Hz): NREMS, F (2, 21) =158.8; Wake, F (2, 21) =13.63. Two-way ANOVA with Tukey’s test in REMS analysis: F (58, 630) =34.56. Data are mean ± s.e.m. * *p*<0.05; ** *p*<0.01; *** *p*<0.001; **** *p*<0.0001. Not significant (*p*>0.05) is not shown.

### *Sik3*^*S551A-L*^ mice exhibit increased NREMS amount with modest effect on NREMS delta power

It would be optimal to compare the sleep phenotypes resulted from isoform-specific expression of SLP-L and SLP-S in the same experimental setting. However, it was impossible to perform ABC-expression of SLP-L because it exceeded the size limit of AAV packaging [20]. Thus, we attempted to construct the *Sik3*^*S551A-L*^ or *Sik3*^*S551A-S*^ knock-in mice by in-frame replacement of *Sik3* exon 13 genomic sequence, respectively, with the C-terminal HA-tagged *Sik3-L* or *Sik3-S* ORF sequence, which carried the S551A mutation and was immediately followed by the bovine growth hormone polyadenylation sequence **(Figure 5, A and B and Supplementary Figure S4A)**. As a result, the *Sik3*^*S551A-L*^ or *Sik3*^*S551A-S*^ mutant allele was expected to express only the long or short isoform of SIK3^S551A^ (SLP). Despite of repeated attempts, we were only able to generate *Sik3*^*S551A-L*^ mice, but not *Sik3*^*S551A-S*^ mice, although, in theory, there should be an equal or better chance of success owning to a smaller size of insert for *Sik3*^*S551A-S*^ mice **(Supplementary Figure S4, A and B)**. We verified that only SIK3^S551A^-L (SLP-L) isoform was expressed in homozygous *Sik3*^*S551A-L*^ mice, whereas the level of *Sik3*^*S551A*^*-L* mRNA was slightly less than that of *Sik3-L* mRNA in heterozygous *Sik3*^*S551A-L*^ mice **(Figure 5, C and D)**. Interestingly, homozygous *Sik3*^*S551A-L*^ mice were born at a sub-mendelian ratio from heterozygote-heterozygote cross, suggesting that isoform-specific expression of SIK3^S551A^-L (SLP-L) might cause early lethality **(Supplementary Figure S4, C and D)**. Likewise, isoform-specific expression of SIK3^S551A^-S (SLP-S) might also result in early lethality and failure to generate *Sik3*^*S551A-S*^ mice.

**Figure 5.**
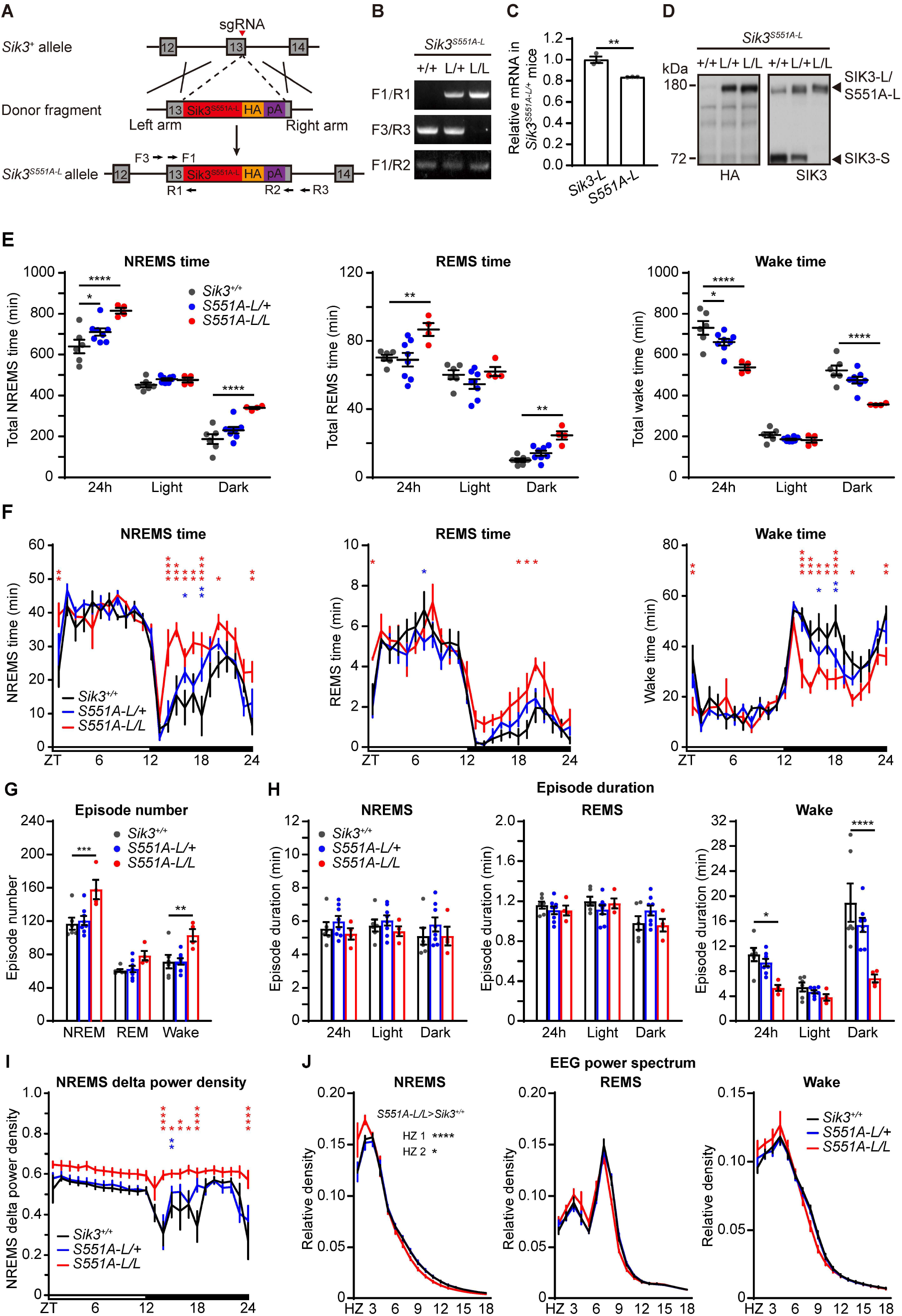
*Sik3*^*S551A-L*^ mice exhibit increased NREMS amount with modest effect on NREMS delta power. **(A)** Schematic for construction of *Sik3*^*S551A-L*^ mice by CRISPR/Cas9 and homologous recombination. **(B)** PCR genotyping of wild-type, heterozygous and homozygous *Sik3*^*S551A-L*^ mice. **(C)** Quantitation of relative expression of *Sik3-L* and *Sik3*^*S551A-L*^ transcripts in heterozygous *Sik3*^*S551A-L*^ mice by RT-qPCR. **(D)** Immunoblotting of whole brain lysates of wild-type, heterozygous and homozygous *Sik3*^*S551A-L*^ mice. **(E)** Quantification of daily NREMS, REMS or Wake time in wild-type (n=6), heterozygous (n=8) and homozygous (n=4) *Sik3*^*S551A-L*^ mice. Two-way ANOVA with Dunnett’s test: NREMS, F (4, 45) =4.259; REMS, F (4, 45) =2.106; Wake, F (4, 45) =5.103. **(F)** Hourly plots of NREMS, REMS or Wake time in wild-type, heterozygous and homozygous *Sik3*^*S551A-L*^ mice. Two-way ANOVA with Dunnett’s test: NREMS, F (46, 360) =1.995; REMS, F (46, 360) =1.041; Wake, F (46, 360) =1.94. **(G and H)** Quantification of NREMS, REMS and Wake episode numbers (G) or episode durations (H) in wild-type, heterozygous and homozygous *Sik3*^*S551A-L*^ mice. Two-way ANOVA with Dunnett’s test: Episode number, F (4, 45) =0.955; NREMS episode duration, F (4, 45) =0.081; REMS episode duration, F (4, 45) =1.837; Wake episode duration, F (4, 45) =3.582. **(I)** Hourly plot of relative NREMS delta power density in wild-type, heterozygous and homozygous *Sik3*^*S551A-L*^ mice. Two-way ANOVA with Dunnett’s test: F (46, 360) =1.181. **(J)** EEG power spectra analysis of NREMS, REMS and Wake states (ZT0-ZT24) in wild-type, heterozygous and homozygous *Sik3*^*S551A-L*^ mice. Two-way ANOVA with Dunnett’s test: NREMS, F (58, 450) =1.824; REMS, F (58, 450) =1.830; Wake, F (58, 450) =1.133. Data are mean ± s.e.m. * *p*<0.05; ** *p*<0.01; *** *p*<0.001; **** *p*<0.0001. Not significant (*p*>0.05) is not shown.

Next, we examined the sleep phenotypes of wild-type, heterozygous and homozygous *Sik3*^*S551A-L*^ mice by EEG/EMG recording. As compared to wild-type littermates, heterozygous *Sik3*^*S551A-L*^ mice showed moderate (average ∼80 min/day) increase of NREMS time, no change in REMS time and decrease of wake time **(Figure 5, E and F)**. Homozygous *Sik3*^*S551A-L*^ mice exhibited marked (average ∼175 min/day) increase of NREMS time, increase of REMS time and decrease of wake time, all of which occurred predominantly during the dark phase **(Figure 5, E and F**). Additionally, *Sik3*^*S551A-L*^ homozygotes showed increased NREMS or wake episode number and shorter wake episode duration during the dark phase **(Figure 5, G and H)**. Whereas wild-type and heterozygous *Sik3*^*S551A-L*^ mice showed similar EEG power spectra profiles, homozygous *Sik3*^*S551A-L*^ mice displayed modest increase of NREMS delta power, particularly in 1-2 Hz frequency range **(Figure 5, I and J)**. Taken together, these results suggest that isoform-specific expression of SLP-L significantly increases NREMS amount with a modest effect on NREMS delta power.

### ABC-expression of SLP-S complements sleepy phenotypes of *Sik3*^*S551A-L*^ mice

A previous study reported that heterozygous *Sik3*^*S551A*^ mice, which expressed all SLP isoforms, exhibited stronger hypersomnia phenotypes than our heterozygous *Sik3*^*S551A-L*^ mice, as shown by average ∼160 min (vs. ∼80 min) increase of daily NREMS amount accompanied with significantly elevated (vs. no change of) NREMS delta power [9]. Thus, we hypothesized that the lack of SLP-S might be responsible for the milder sleep phenotypes of *Sik3*^*S551A-L*^ mice. To test this possibility, we performed ABC-expression of SLP-S in the adult brain neurons of heterozygous *Sik3*^*S551A-L*^ mice by intravenous injection of AAV-hSyn-mCherry or AAV-hSyn-SLP-S **(Figure 6A)**. Accordingly, ABC-expression of SLP-S, relative to ABC-expression of mCherry, further increased NREMS amount and NREMS delta power in *Sik3*^*S551A-L*^ mice to levels approaching those of *Sik3*^*S551A*^ or *Sik3*^*Slp*^ mice **(Figure 6, B-G)** [1,9]. While there was increased REMS time in heterozygous *Sik3*^*S551A-L*^ mice, ABC-expression of SLP-S reversed this phenotype by moderately reducing REMS amount during the light phase, which was consistent with the REMS phenotype of *Sik3*^*S551A*^ or *Sik3*^*Slp*^ mice [1,9]. Collectively, these results indicate that both long and short isoforms of SLP kinase contribute critically to the hypersomnia phenotypes of *Sik3*^*S551A*^ or *Sik3*^*Slp*^ mice.

**Figure 6.**
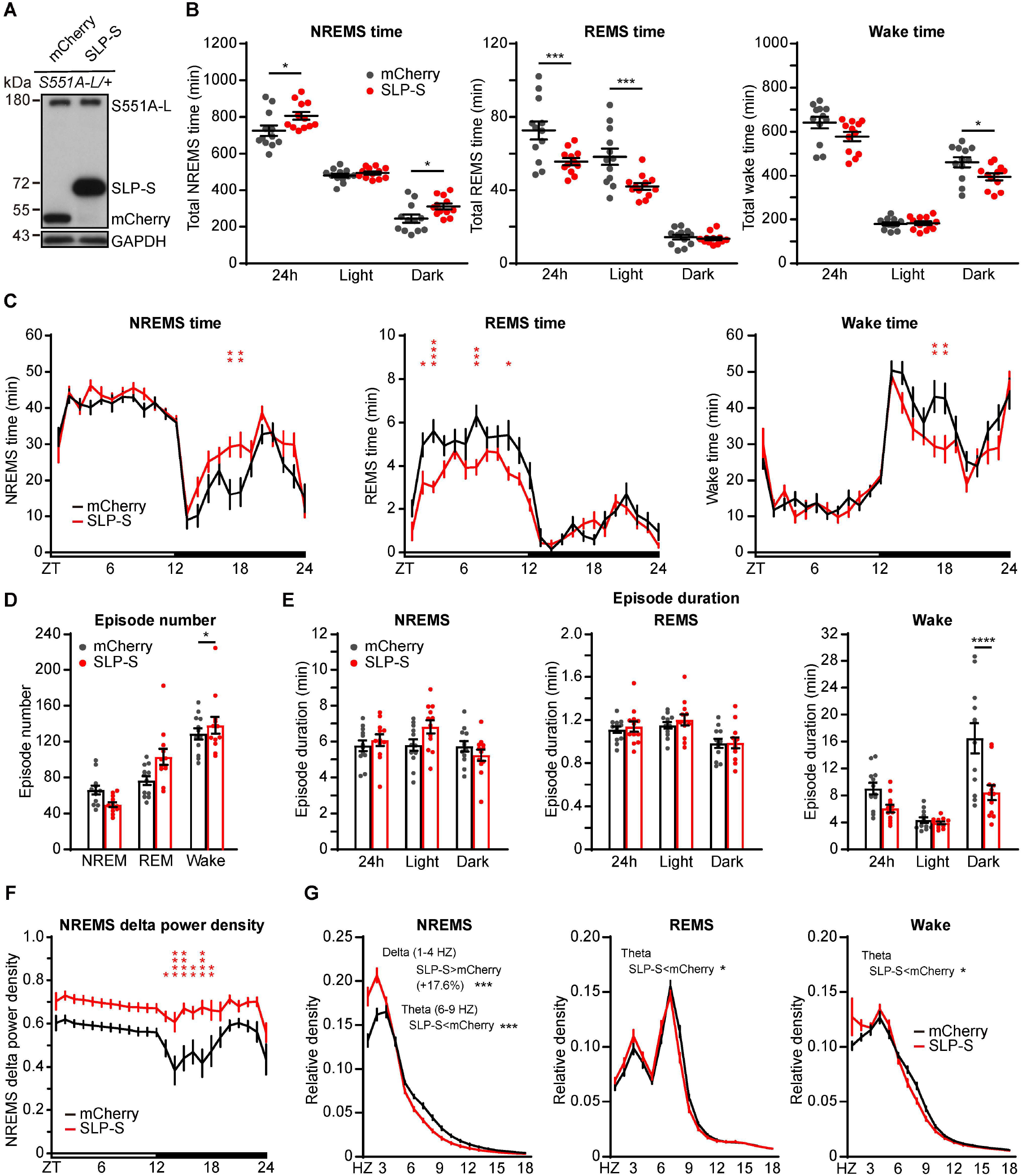
ABC-expression of SLP-S complements sleep phenotypes in *Sik3*^*S551A-L*/+^ mice. **(A)** Immunoblotting of whole brain lysates prepared from AAV-hSyn-mCherry or AAV-hSyn-SLP-S injected heterozygous *Sik3*^*S551A-L*^ mice. **(B)** Quantification of daily NREMS, REMS or Wake time in the AAV-hSyn-mCherry (n=12) or AAV-hSyn-SLP-S (n=12) injected *Sik3*^*S551A-L*/+^ mice. Two-way ANOVA with Sidak’s test: NREMS, F (2, 66) =1.724; REMS, F (2, 66) =4.63; Wake, F (2, 66) =2.223. **(C)** Hourly plots of NREMS, REMS or Wake time of the ABC-mCherry or ABC-SLP-S *Sik3*^*S551A-L*/+^ mice. Two-way ANOVA with Sidak’s test: NREMS, F (23, 528) =1.619; REMS, F (23, 528) =2.584; Wake, F (23, 528) =1.884. **(D and E)** Quantification of NREMS, REMS and Wake episode numbers (D) or episode durations (E) of the ABC-mCherry or ABC-SLP-S *Sik3*^*S551A-L*/+^ mice. Two-way ANOVA with Sidak’s test: Episode number, F (2, 66) =5.253; NREMS episode duration, F (2, 66) =2.838; REMS episode duration, F (2, 66) =0.141; Wake episode duration, F (2, 66) =6.067. **(F)** Hourly plot of relative NREMS delta power density in the ABC-mCherry or ABC-SLP-S *Sik3*^*S551A-L*/+^ mice. Two-way ANOVA with Sidak’s test. F (23, 528) =0.979. **(G)** EEG power spectra analysis of NREMS, REMS and Wake states (ZT0-ZT24) in the ABC-mCherry or ABC-SLP-S *Sik3*^*S551A-L*/+^ mice. Unpaired t test: Delta (1-4 Hz): NREMS, t =4.071; REMS, t =1.758; Wake, t =1.795. Theta (6-9 Hz): NREMS, t =4.626; REMS, t =2.208; Wake, t =2.487. df =22. Data are mean ± s.e.m. * *p*<0.05; ** *p*<0.01; *** *p*<0.001; **** *p*<0.0001. Not significant (*p*>0.05) is not shown.

## DISCUSSION

In this study, we identified a most abundant short isoform of SIK3/SLP and characterized the sleep-promoting activity of SLP-S in adult mice by AAV-mediated somatic genetics analysis [12]. ABC-expression of SLP-S or SLP^T221E^-S, which carried the kinase-activating mutation, in the adult brain neurons caused modest (∼50 min/day) or marked (∼145 min/day) increase of NREMS amount, respectively, accompanied with significantly and constitutively elevated NREMS delta power. By contrast, ABC-expression of SLP^K37M^-S or SLP^T221A^-S, which carried kinase-inactivating mutations [11,15], resulted in little or no sleep phenotype, suggesting that the sleep-promoting activity of SLP-S is dependent on its kinase activity.

We also examined the sleep phenotypes resulted from isoform-specific expression of SLP-L by constructing the *Sik3*^*S551A-L*^ knock-in mice. Heterozygous *Sik3*^*S551A-L*^ mice displayed much less NREMS amount and lower NREMS delta power than heterozygous *Sik3*^*S551A*^ mice [9]. Moreover, ABC-expression of SLP-S complemented sleep phenotypes of *Sik3*^*S551A-L/+*^ mice by further increasing NREMS amount and NREMS delta power to levels of *Sik3*^*S551A/+*^ mice [9]. These gain-of-function experiments indicate that both SLP-L and SLP-S play important roles in the regulation of sleep quantity and intensity in *Sik3*^*Slp*^ or *Sik3*^*S551A*^ mice. Consistently, partial loss-of-function of *Sik3* reduced NREMS delta power and impaired homeostatic response to sleep deprivation [15]. In the absence of isoform-specific knockout studies, however, it is difficult to draw firm conclusions about the normal functions of SIK3-L and SIK3-S in sleep regulation.

We observed significantly increased daily NREMS amount accompanied with modest elevation of NREMS delta power in homozygous *Sik3*^*S551A-L*^ mice. On the other hand, ABC-expression of SLP-S in the adult brain neurons significantly elevated NREMS delta power, whereas slightly increases NREMS amount. These observations suggest that NREMS amount is not strictly linked with NREMS delta power, which may be regulated separately. Moreover, *Sik3*^*S551A-L*^ mice, which lacked the expression of SLP-S, increased REMS amount during the dark phase, whereas *Sik3*^*S551A*^ mice showed reduction of REMS amount during the light phase. Accordingly, ABC-expression of SLP-S caused reduction of REMS time during the light phase, suggesting that SLP-S, rather than SLP-L, contributed to reduction of REMS amount in *Sik3*^*Slp*^ or *Sik3*^*S551A-L*^ mice. Whereas expression of SIK3^S551A^-L was comparable to that of SIK3-L in *Sik3*^*S551A-L/+*^ mice, AAV-mediated expression of SLP-S was significantly less than that of endogenous SIK3-S in wild-type or *Sik3*^*S551A-L/+*^ mice. These observations argue against a simple explanation of dosage effect for the different sleep phenotypes resulted from isoform-specific expression SLP-L and SLP-S. Taken together, these results suggest the possibility that SLP-L and SLP-S may have quantitatively different effects on sleep quantity and intensity.

Recent studies have shown that SLP/SIK3 signaling in distinct populations of excitatory neurons of the hypothalamus and cortex regulates NREMS amount and NREMS delta power, respectively (Yanagisawa M. and Funato H., unpublished). Interestingly, the relative expression of SIK3-S vs. SIK3-L/M proteins varied greatly among different brain regions, with a high ratio of 57.3:1 in the cortex and a low ratio of 5.5:1 in the hypothalamus. Thus, it is plausible that SIK3/SLP isoforms may contribute distinctly to regulation of sleep quantity and intensity in different brain regions. Based on our results, we propose that SLP-S may have a prominent effect on NREMS delta power in cortical neurons, whereas both SLP-L and SLP-S may regulate NREMS amount in hypothalamic neurons.

Because SLP-S lacks the C-terminal half portion of SLP-L, it is interesting to speculate that SLP-L and SLP-S may have different biochemical properties, e.g., interacting with different cofactors or phosphorylating a distinct subset of substrate proteins. By quantitative phosphoproteomic analysis of *Sleepy* mutant and sleep-deprived wild-type mouse brains, we’ve previously identified 80 mostly synaptic sleep need index phosphoproteins (SNIPPs) [14]. Preferential association between SNIPPs and SLP (relative to SIK3) suggest that specific SNIPPs may function as downstream substrates of SLP/SIK3 kinase that mediate sleep regulation [14]. Other known substrates of SIK3 include transcriptional modulators, such as class IIa histone deacetylases (HDACs) and CREB-regulated transcription coactivators (CRTCs)/transducers of regulated CREB activity (TORCs) [21,22]. Therefore, future studies are warranted to identify isoform-specific binding proteins or substrates of SLP/SIK3 by immunoprecipitation and mass spectrometric analysis, which may provide new insights into the molecular mechanisms by which sleep quantity and intensity is regulated in mammals.

It should also be noted that the current study had several inherent limitations, such as the exclusive use of gain-of-function approach, the lack of spatial control of AAV expression by retro-orbital injection, the lack of investigation in female mice, and inability to compare sleep phenotypes resulted from expression of SLP-L and SLP-S in the same experimental setting for technical reasons.

## Acknowledgements

We thank Drs. H. Funato, M. Yanagisawa and A. Ikkyu for useful discussion; K. Wu, W. Min, Y. Zhuang for technical assistance; and Dr. Q. Li for comments on the manuscript. This study was supported by the National Major Project of China Science and Technology Innovation 2030 for Brain Science and Brain-Inspired Technology (2021ZD0203400 to Q.L.); the Beijing Municipal Science and Technology Commission (Z181100001318004 to Q.L.) and the National Key Research and Development Program of China.

## List of Abbreviations

(AAV): Adeno-associated virus
(ABC): Adult brain chimeric
(SIK3): salt-inducible kinase 3
(SLP): SLEEPY
(AMPK): AMP-activated protein kinase
(NREMS): non-rapid eye movement sleep
(REMS): rapid eye movement sleep
(EEG): electroencephalogram
(EMG): electromyogram
(SNIPP): sleep need index phosphoprotein
(HDAC): histone deacetylase
(CRTC): CREB-regulated transcription coactivator
(TORC): transducers of regulated CREB activity
(RT-PCR): reverse transcription and PCR
(SD6): 6-h sleep deprivation
(S6): 6-h ad libitum sleep
(RT): room temperature
(ORF): open reading frame
(PEI): polyethylenimine
(ZT): zeitgeber time
(BCA): bicinchoninic acid
(PBS): phosphate-buffered saline
(GFAP): glial fibrillary acidic protein
(hSyn): human synapsin

