## Supplementary figure legends for "Regulation of sleep quantity and intensity by long and short isoforms of SLEEPY kinase"

^1^College of Life Sciences, Beijing Normal University, Beijing 100875, China; ^2^National Institute of Biological Sciences (NIBS), Beijing 102206, China; ^3^College of Biological Sciences, China Agriculture University, Beijing 100094, China; ^4^Graduate School of Peking Union Medical College, Chinese Academy of Medical Sciences, Beijing 100730, China; ^5^International Institute for Integrative Sleep Medicine (WPI-IIIS), University of Tsukuba, Tsukuba 305-8575, Japan; ^6^Tsinghua Institute of Multidisciplinary Biomedical Research, Tsinghua University, Beijing 102206, China

^7^These authors contributed equally to this study

* Corresponding authors:

Qinghua Liu,;

National Institute of Biological Sciences, Beijing (NIBS)

7 Science Park Rd., ZGC Life Science Park

Beijing, China 102206

**Supplementary Figures**

**Figure S1 (linked to Figure 1) Identification and characterization of a new SIK3/SLP-S isoform.**

**(A)** Immunoblotting of whole brain lysates from wild-type and ABC-*Sik3^KO^* mice by CRISPR/Cas9 [12] using anti-SIK3 antibodies. **(B)** Alignment of the actual and annotated (UniProt F6U8X4) SIK3-S protein sequences. Highlighted sequences include the N-terminal 58 amino acids (blue), protein kinase domain (gray), K37 (red) and exon 13-encoded 52 amino acids (orange) that are deleted in SLP-S. **(C)** Immunoprecipitation of FLAG-tagged SIK3-L, SLP-L or SLP-S from transfected 293T cells and followed by immunoblotting using anti-FLAG and anti-14-3-3 antibodies.

**Figure S2 (linked to Figure 2) ABC-expression of SIK3 or SLP-S in neurons and astrocytes.**

**(A)** Representative images showing co-immunostaining of HA (red) and NeuN (green) in the sagittal brain sections of the AAV-hSyn-GFP, AAV-hSyn-SLP-S or AAV-hSyn-SIK3-S injected C57BL/6J mice. Scale bars, 1mm. **(B-D)** Representative images showing co-immunostaining of HA (red) and NeuN (green) and quantification of the percentage of HA^+^/NeuN^+^ neurons in the cortex (B), thalamus (C) or hypothalamus (D) of the ABC-GFP, ABC-SLP-S or ABC-SIK3-S mice. Scale bars, 20μm. **(E)** Immunoblotting of whole brain lysates of ABC-GFP, ABC-SLP-S or ABC-SIK3-S mice. **(F)** Quantitation of relative expression of *Slp-S* vs. endogenous *Sik3-S* transcripts in ABC-SLP-S mice by RT-qPCR. Data are mean ± s.e.m. Unpaired t test, t =0.0089, df =4, Not significant (*p*>0.05) is not shown. **(G)** Immunoblotting of whole brain lysates of ABC-GFP and ABC-SLP-S mice. **(H)** Quantitation of relative expression of SLP-S vs. endogenous SIK-S proteins in ABC-SLP-S mice (n=3). Data are mean ± s.e.m. Unpaired t test, t =7.874, df =4, ** *p*<0.01. **(I)** Representative images showing co-immunostaining of HA (red) and S100β or NeuN (green) and quantification of the AAV transduction rates of astrocytes and neurons in the cortex of the AAV-GFAP-GFP or AAV-GFAP-SLP-S injected C57BL/6J mice. Scale bars, 20μm.

**Figure S3 (linked to Figure 2) ABC-expression of SLP-S in astrocytes does not cause sleepy phenotypes.**

**(A)** Quantification of daily NREMS, REMS or Wake time in the AAV-GFAP-GFP (n=10) and AAV-GFAP-SLP-S (n=8) injected mice. Two-way ANOVA with Sidak's test: NREMS, F (2, 48) =2.109; REMS, F (2, 48) =0.075; Wake, F (2, 48) =1.818. **(B)** Hourly plots of NREMS, REMS or Wake time in the AAV-GFAP-GFP and AAV-GFAP-SLP-S injected mice. Two-way ANOVA with Sidak's test: NREMS, F (23, 384) =2.517; REMS, F (23, 384) =0.702; Wake, F (23, 384) =2.219. **(C and D)** Quantification of NREMS, REMS and Wake episode numbers (C) and episode durations (D) of the AAV-GFAP-GFP and AAV-GFAP-SLP-S injected mice. Two-way ANOVA with Sidak's test: Episode number, F (2, 48) =3.959; NREMS episode duration, F (2, 48) =7.415; REMS episode duration, F (2, 48) =0.101; Wake episode duration, F (2, 48) =1.067. **(E)** Hourly plot of relative NREMS delta power density in the AAV-GFAP-GFP and AAV-GFAP-SLP-S injected mice. Two-way ANOVA with Sidak's test: F (23, 384) =2.475. **(F)** EEG power spectra analysis of NREMS, REMS and Wake states (ZT0-ZT24) in the AAV-GFAP-GFP and AAV-GFAP-SLP-S injected mice. Unpaired t test. Delta (1-4 Hz): NREMS, t =4.018; REMS, t =3.989; Wake, t =2.787. Theta (6-9 Hz): NREMS, t =5.068; REMS, t =5.117; Wake, t =2.379. df =16. Data are mean ± s.e.m. * *p*<0.05; ** *p*<0.01; *** *p*<0.001; **** *p*<0.0001. Not significant (*p*>0.05) is not shown.

**Figure S4 (linked to Figure 5) Generation of homozygous *Sik3^S551A-L^* mice.**

**(A)** Schematic for the sgRNA target sequence and donor fragments for generating *Sik3^S551A-L^* or *Sik3^S551A-S^* knock-in mice by CRISPR/Cas9 and homologous recombination. **(B)** The number of injected zygotes and percentage of positive founders for *Sik3^S551A-L^* or *Sik3^S551A-S^* mice. **(C)** The number and percentage of wild-type, heterozygous and homozygous *Sik3^S551A-L^* progenies were generated from the heterozygote-heterozygote mating. **(D)** Statistic analysis of the deviations from Mendelian genetics of the heterozygote-heterozygote mating.
