## Supplementary figures and images for "Regulation of sleep quantity and intensity by long and short isoforms of SLEEPY kinase"

### Supplementary figure 1

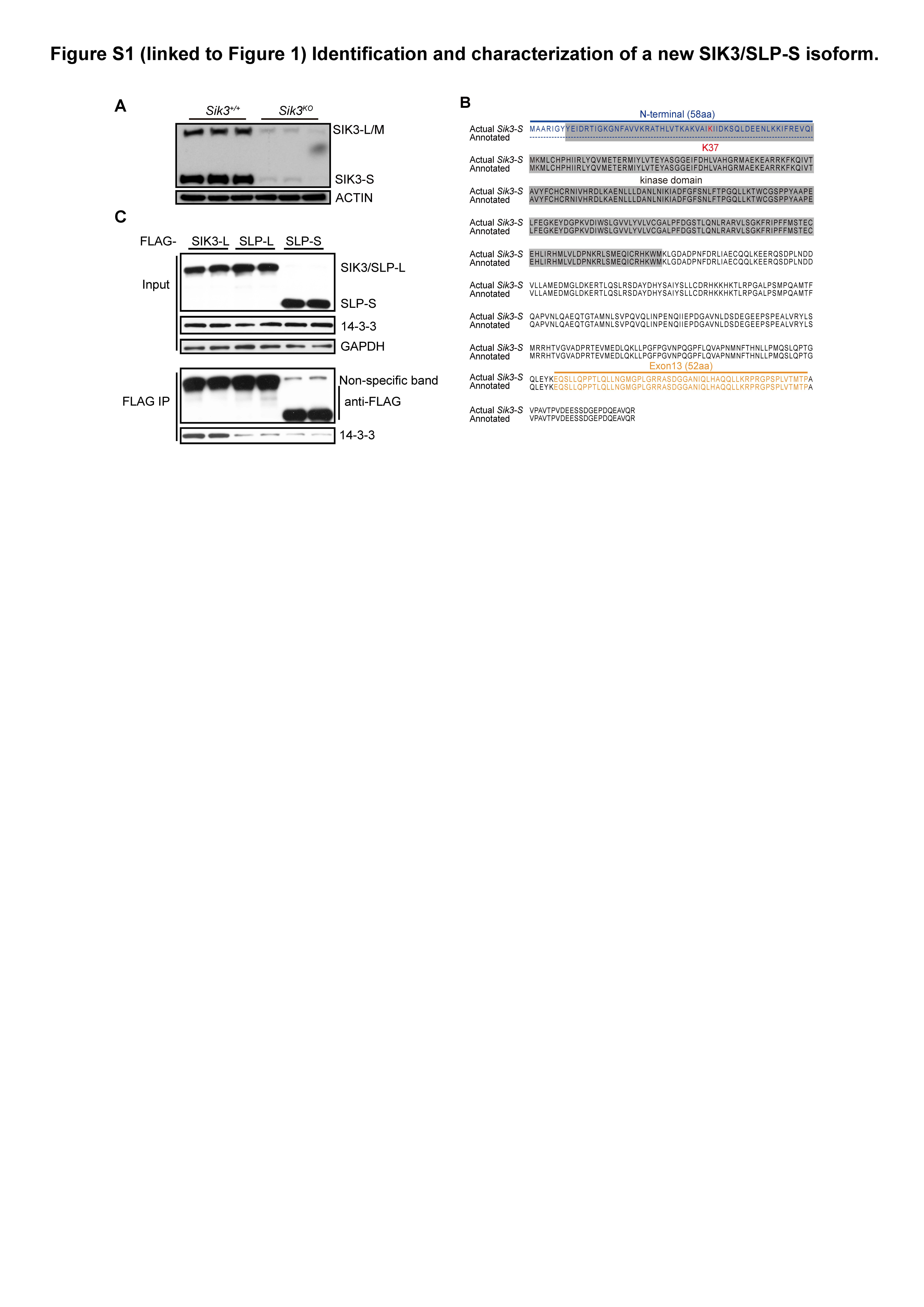

### Supplementary figure 2

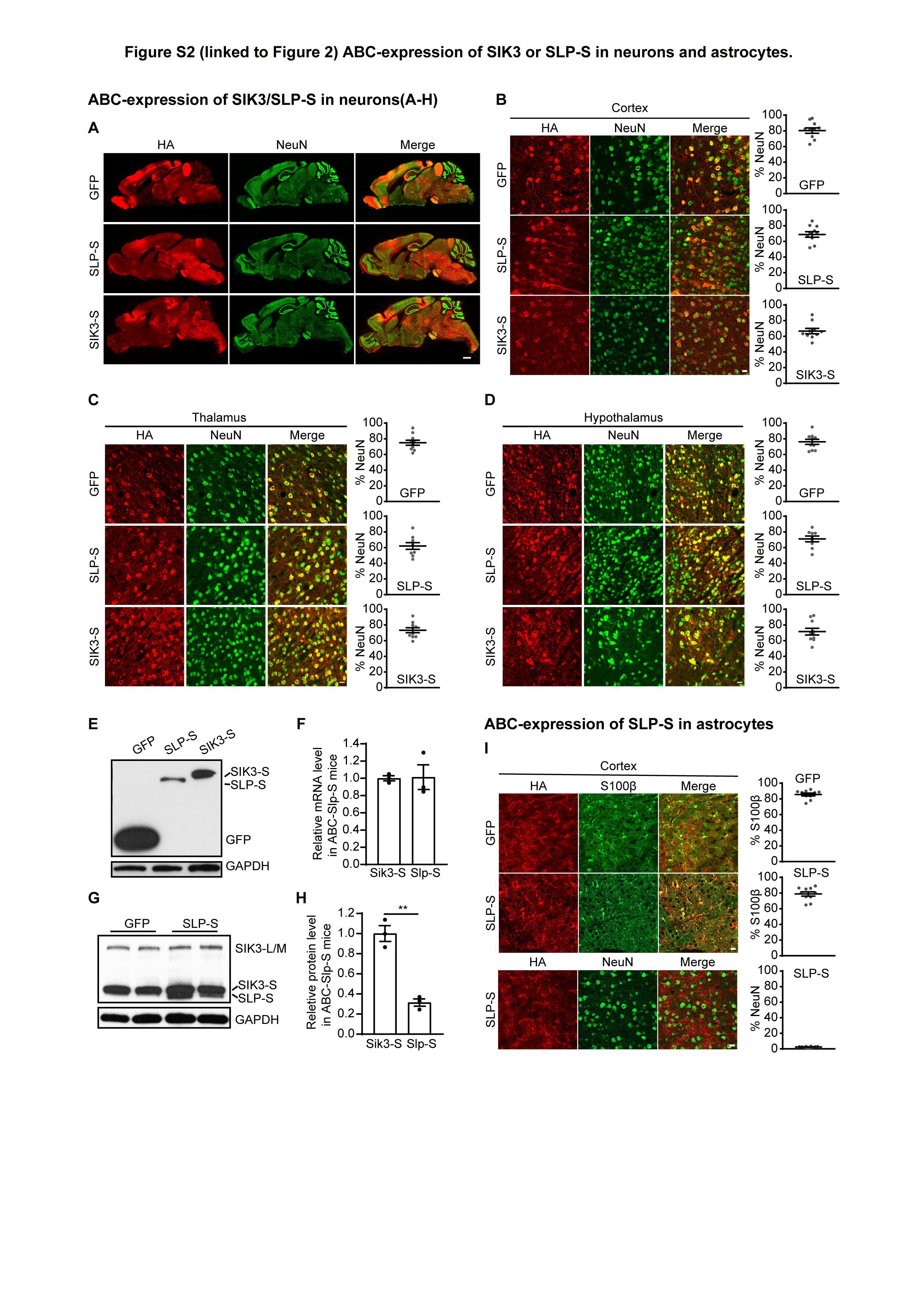

### Supplementary figure 3

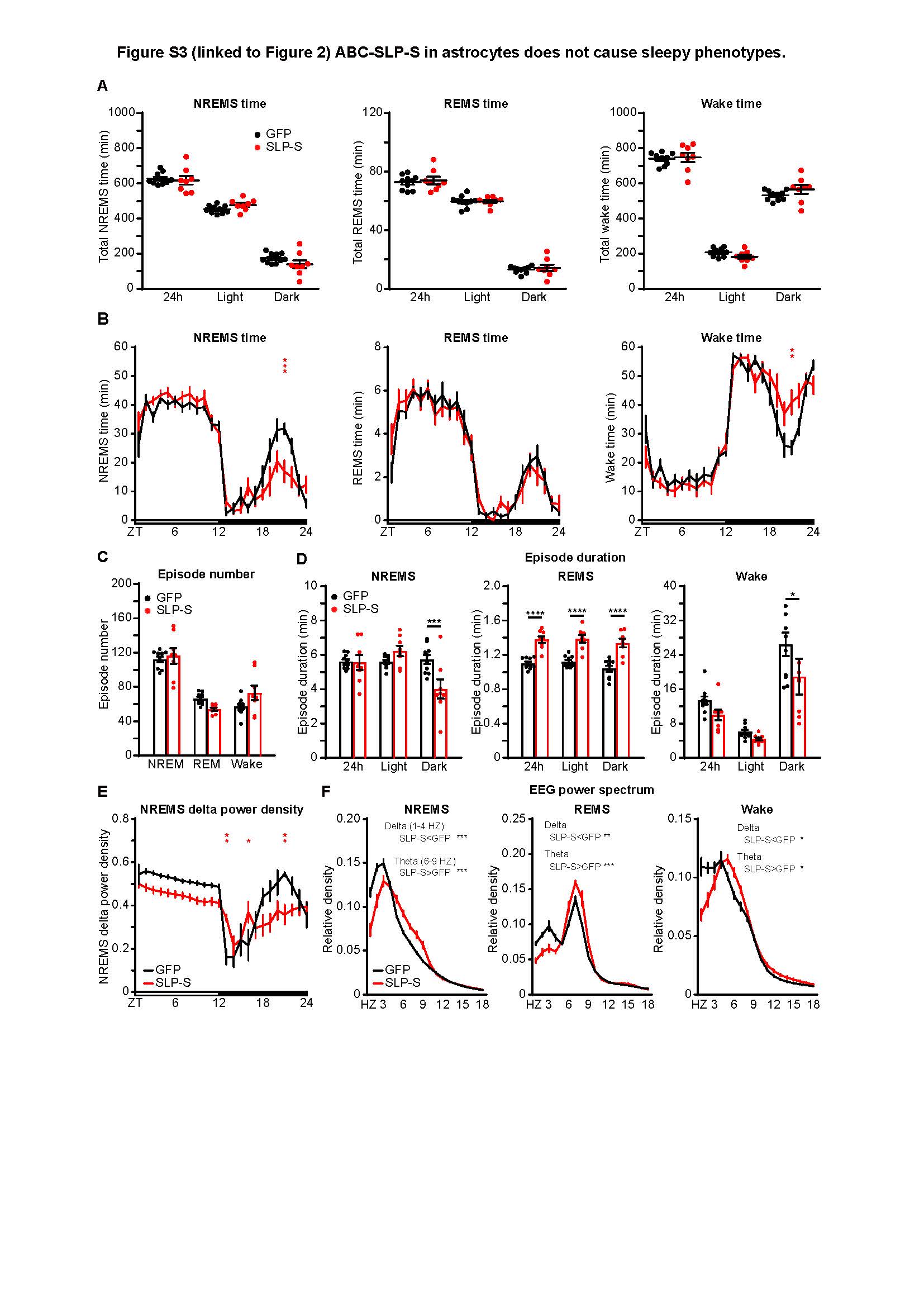

### Supplementary figure 4

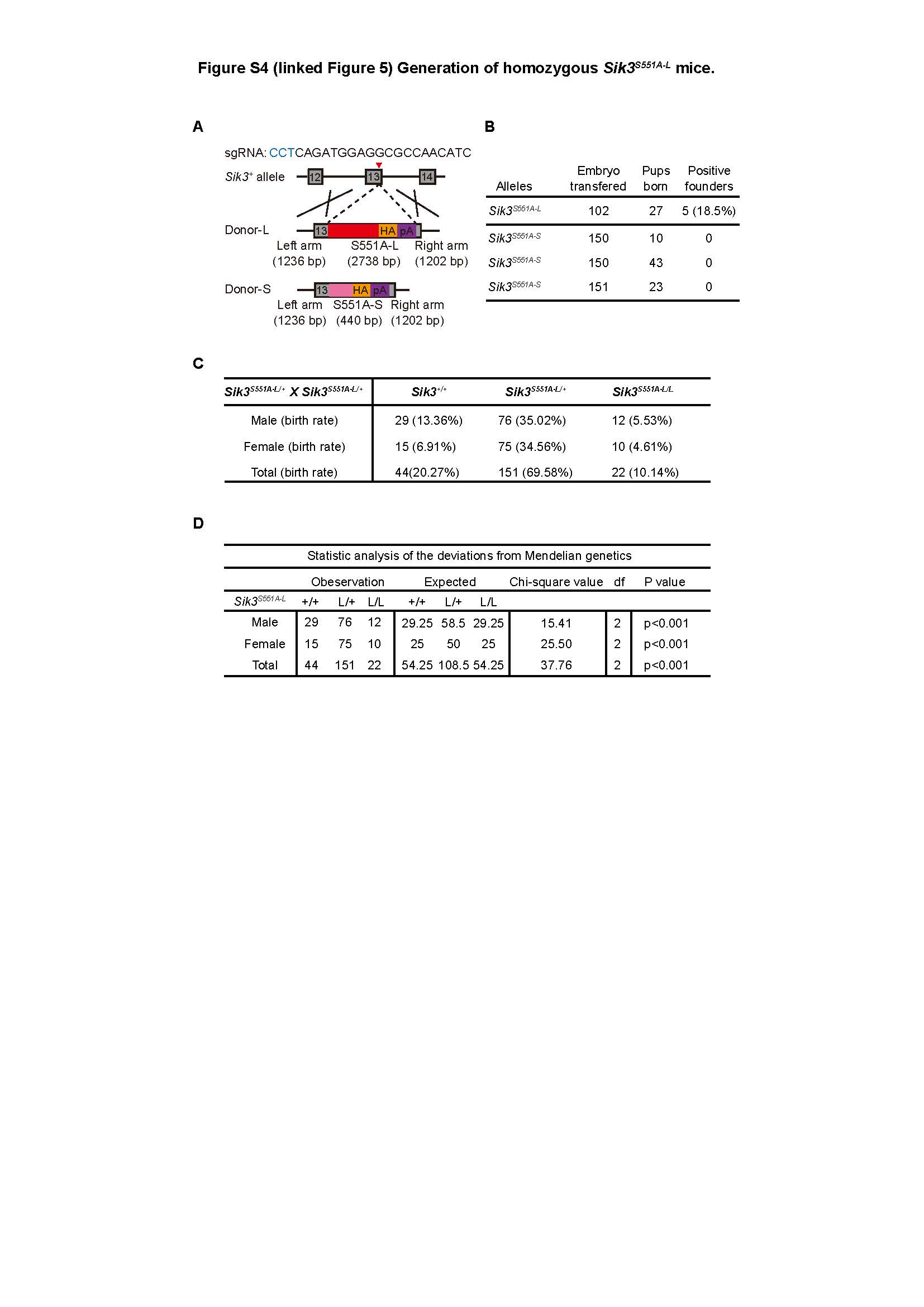
